# CIP2A Mediates the Recruitment of the SLX4-MUS81-XPF Tri-Nuclease Complex in Mitosis and Protects Against Replication Stress

**DOI:** 10.1101/2025.09.19.677284

**Authors:** Alice Meroni, Annica Pellizzari, Andrea Hänel, Nathalie Varisco, Pascale Brasier-Lutz, Isabell Witzel, Alessandro A Sartori, Manuel Stucki

## Abstract

DNA replication stress frequently elicits problems in mitosis because incompletely replicated chromosomes or replication intermediates physically link sister chromatids together and prevent their proper segregation during anaphase. We and others recently discovered a mitotic role for the CIP2A oncoprotein, which is critically implicated in chromosome stability maintenance and chromosome fragment clustering during mitosis. In addition, in homologous recombination deficient (HRD) cells, CIP2A is essential and may thus constitute a new drug target in HRD cancers. Yet, the precise mechanisms by which CIP2A suppresses chromosomal instability during mitosis and thus allows for the survival of HRD cancer cells remain largely elusive. Here we characterize CIP2A’s role in DNA replication stress responses. We show that upon replication stress, wild-type cells show an elevated accumulation of CIP2A foci during mitosis, indicating its involvement in the mitotic response to replication stress. Defective DNA replication leads to the accumulation of under-replicated DNA, which can be carried into mitosis. We demonstrate that in the absence of CIP2A, cells fail to recruit the SLX4-MUS81-XPF (SMX) tri-nuclease complex to sites of under-replicated DNA in mitosis, resulting in a high incidence of lagging chromosomes during anaphase and subsequent micronuclei formation. In a subset of cell lines, we also observed CIP2A-dependent mitotic DNA synthesis (MiDAS) upon replication stress. However, our data suggest that MiDAS and SMX recruitment are not functionally linked. This novel role of CIP2A in managing under-replicated DNA may provide insights into the molecular mechanisms underlying therapeutic vulnerabilities in cancer cells.

## Introduction

Faithful DNA replication is essential for genome stability, yet replication stress – arising from obstacles such as DNA damage, replication fork stalling, and oncogene activation – poses a significant threat to genomic integrity (Zeman & Cimprich, 2014). When replication is challenged during the S-phase, cells may enter mitosis with under-replicated DNA and single strand DNA gaps. In this state, the sister chromatids could remain physically linked by Watson-Crick base pairing, which can hinder their proper segregation during anaphase. To prevent catastrophic chromosomal segregation errors, cells activate mitotic DNA synthesis (MiDAS), an atypical replication mechanism specifically activated in early mitosis to resolve these under-replicated DNA regions and ensure proper chromosome segregation (Bhowmick et al., 2023).

MiDAS is primarily observed under conditions of replication stress, particularly at late-replicating genomic regions such as common fragile sites (CFSs) (Le Beau et al., 1998). In most of the studies, MiDAS is typically induced by low doses of the DNA polymerase inhibitor aphidicolin (Aph), which slows replication and triggers transcription-replication conflicts at CFSs (Glover et al., 1984). Mechanistically, MiDAS involves a series of coordinated actions of nucleases and polymerases to complete DNA synthesis that could resemble break-induced replication (BIR). Under-replicated DNA sites are first marked by FANCD2 (Chan et al., 2009), followed by the recruitment of TOPBP1 (Pedersen et al., 2015; Leimbacher et al., 2019; Gallina et al., 2016), which facilitates the assembly of additional factors. Among these factors, SLX4 is a crucial scaffold protein that coordinates recruitment and activation of structure-specific nucleases such as MUS81-EME1 (Minocherhomji et al., 2015; Pedersen et al., 2015). In addition to nuclease activity, MiDAS also requires factors involved in DNA synthesis and repair, such as the strand-annealing protein RAD52 (Bhowmick et al., 2016), the POLD3 subunit of polymerase δ (Bhowmick et al., 2016), and translesion synthesis polymerases Pol ζ and REV1 (Wu et al., 2023). However, the full regulatory network governing mitotic responses to replication stress remains unclear, and the precise coordination of these factors is yet to be fully elucidated. Identifying novel regulators involved in this process could help clarify mechanisms of genome stability maintenance and reveal vulnerabilities exploitable in cancer therapy.

CIP2A (<u>C</u>ancerous Inhibitor of <u>p</u>rotein phosphatase <u>2A</u>) is an oncogenic factor initially characterized as an inhibitor of protein phosphatase 2A (PP2A), involved in promoting tumor growth and proliferation (Junttila et al., 2007). Recently, CIP2A has been implicated in protecting chromosome integrity during mitosis (De Marco Zompit & Stucki, 2021; Adam et al., 2021). CIP2A interacts with Topoisomerase II-binding protein 1 (TOPBP1), forming a complex essential for tethering DNA double-strand breaks (DSBs) during mitosis in an MDC1-dependent fashion (De Marco Zompit et al., 2022). Intriguingly, CIP2A has also emerged as a synthetic lethal partner of BRCA1 and BRCA2, highlighting its potential as a therapeutic target in cancers harboring these mutations. However, the mechanistic basis of this interaction remains unclear (Adam et al., 2021) and defining how CIP2A functions within this context may offer valuable therapeutic insights.

Notably, cells deficient in BRCA2 spontaneously activate MiDAS, likely as a compensatory response to chronic replication stress (Feng & Jasin, 2017; Groelly et al., 2022). Given that CIP2A exhibits synthetic lethality with BRCA deficiency, and it directly interacts with TOPBP1 in mitosis, we hypothesize that CIP2A might retain a specific function in mitigating replication stress during mitosis.

Here, we investigate the role of CIP2A in mitotic replication stress response after loss of BRCA2, upon ATR inhibition, and in response to DNA replication inhibition. We demonstrate that CIP2A regulates the recruitment of the structure-specific endonuclease complex SLX4-MUS81-XPF (SMX complex), and that its loss leads to increased chromosomal instability under replication stress, manifesting as anaphase defects and micronuclei formation. Our data further indicate that in U2OS, but not in DLD1 cells, MiDAS is dependent on CIP2A upon replication stress. Importantly, our data indicate that during mitosis, the recruitment of the SMX complex in microscopically discernible foci and MiDAS may be genetically separable processes that are not functionally linked. Altogether, we provide mechanistic insights into how mitotic replication stress is resolved to maintain genome stability.

## Results and Discussion

### The CIP2A-TOPBP1 Complex Responds to Replication Stress in Mitosis

To investigate CIP2A’s role in response to replication stress, we first looked at its recruitment in the form of foci during mitosis under various replication stress conditions. Consistent with our previous findings (Adam et al., 2021), CIP2A and TOPBP1 foci were significantly increased in BRCA2 KO DLD1 mitotic cells (Fig. 1A, B). In addition, the proximity of CIP2A with TOPBP1 was enhanced specifically in BRCA2 KO mitotic cells, as determined by proximity ligation assay (PLA; Fig. 1C).

**Figure 1:**
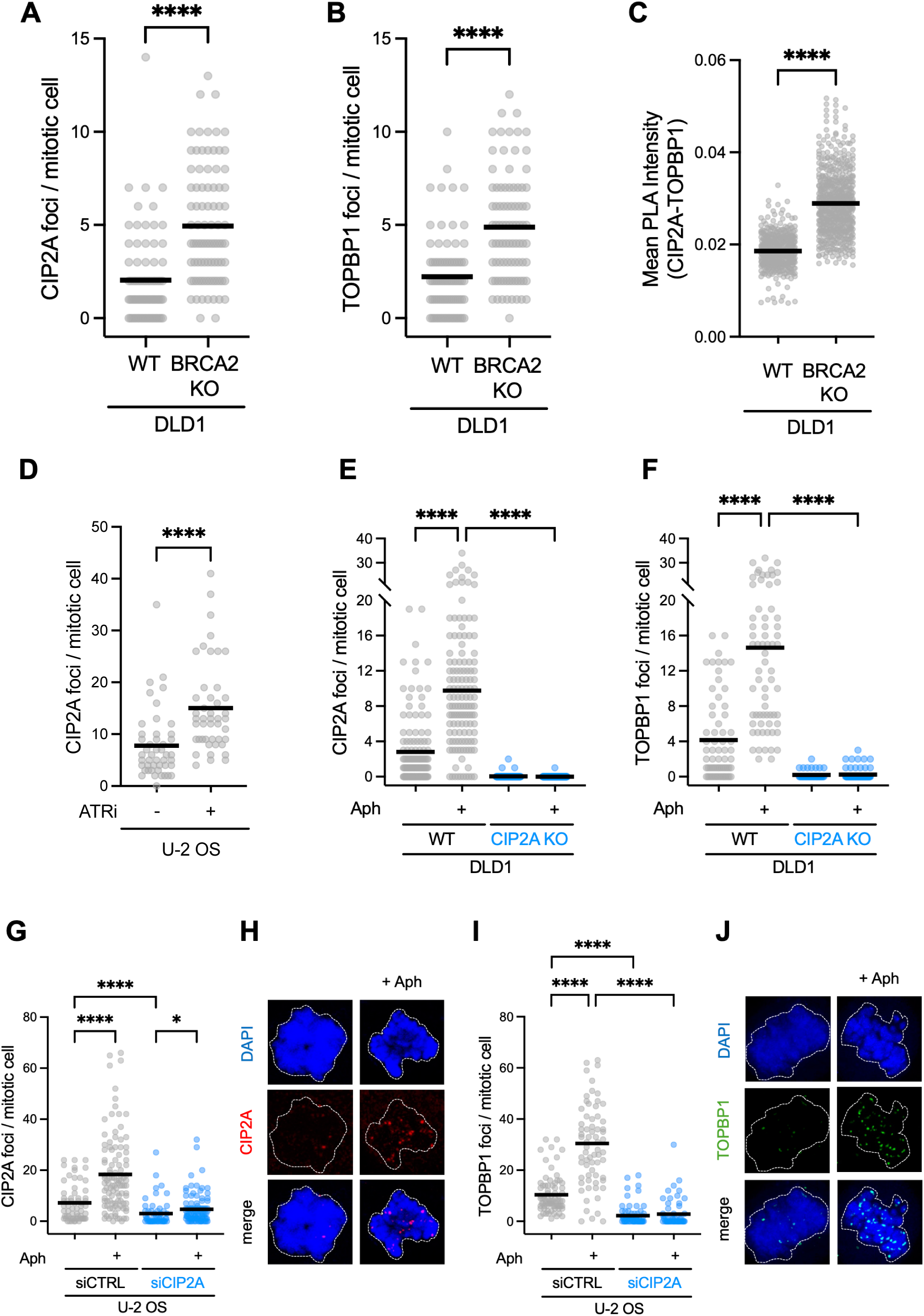
The CIP2A-TOPBP1 complex responds to replication stress in mitosis. **A)** Dot plot showing CIP2A foci per mitotic cells in DLD1 WT and DLD1 BRCA2 knockout cells. Statistical analysis: Mann-Whitney test, n = 2 independent replicates. **B)** Dot plot showing TOPBP1 foci per mitotic cells in DLD1 WT and DLD1 BRCA2 knockout cells. Statistical analysis: Mann-Whitney test, n = 2 independent replicates. **C)** Quantitative analysis of the mean proximity ligation assay (PLA) intensity in DLD1 WT and DLD1 BRCA2 knockout cells stained for CIP2A and TOPBP1. Statistical analysis: Welch’s t-test. n = 2 independent replicates. **D)** Dot plot showing CIP2A foci per mitotic cell in U2OS cells treated with ATR inhibitor (ATRi, 10 μM) for 24 h following siRNA depletion. Cells were synchronized in RO-3306 (7 μM) for 6 h. Mean indicated. Statistical analysis: Kruskal-Wallis test, followed by uncorrected Dunn’s multiple comparisons test. n = 2 independent replicates. **E)** Dot plot showing CIP2A foci per mitotic cell in DLD1 cells treated with Aph (0.4 μM) for 24 h. Cells were synchronized in RO-3306 (7 μM) for 7 h. Median indicated. Statistical analysis: Kruskal-Wallis test, followed by uncorrected Dunn’s multiple comparisons test. n = 4 independent replicates. **F)** Dot plot showing TOPBP1 foci per mitotic cell in DLD1 cells treated with Aph (0.4 μM) for 24 h. Cells were synchronized in RO-3306 (7 μM) for 7 h. Median indicated. Statistical analysis: Kruskal-Wallis test, followed by uncorrected Dunn’s multiple comparisons test. n = 3 independent replicates. **G)** Dot plot showing CIP2A foci per mitotic cell in U2OS cells treated with Aph (0.4 μM) for 24 h following siRNA depletion. Cells were synchronized in RO-3306 (7 μM) for 7 h. Median indicated. Statistical analysis: Kruskal-Wallis test, followed by uncorrected Dunn’s multiple comparisons test. n = 3 independent replicates. **H)** Confocal microscopy images showing CIP2A foci (red) in mitotic U2OS cells, with nuclei stained with DAPI (blue). Images are maximum projections of Z-stacks. **I)** Dot plot showing TOPBP1 foci per mitotic cell in U2OS cells treated with Aph (0.4 μM) for 24 h following siRNA depletion. Cells were synchronized in RO-3306 (7 μM) for 7 h. Median indicated. Statistical analysis: Kruskal-Wallis test, followed by uncorrected Dunn’s multiple comparisons test. n = 2 independent replicates. **J)** Confocal microscopy images showing TOPBP1 foci (green) in mitotic U2OS cells, with nuclei stained with DAPI (blue). Images are maximum projections of Z-stacks.

Next, we induced replication stress by inhibiting ATR with VE-821 (ATRi) and found that CIP2A foci are increased in mitotic cells (Fig. 1D), suggesting that CIP2A not only responds to BRCA-deficiency-associated replication stress but also to ATRi-induced replication stress. Notably, CIP2A-deficient cells exhibit high sensitivity to ATR inhibitor treatment, but the underlying cause remains unknown (Hustedt et al., 2019).

Finally, we induced replication stress using low doses of Aph, a replicative DNA polymerase inhibitor, to further assess whether CIP2A-TOPBP1 responds to stalled replication forks and under-replicated DNA in mitosis. Consistently, CIP2A and TOPBP1 were recruited in response to Aph-induced replication stress in mitotic cells, as observed in both DLD1 (Fig. 1E, F) and U2OS cells (Fig. 1G, H, I, J). Importantly, TOPBP1 recruitment was abolished in the absence of CIP2A, confirming the co-dependency of these two factors (De Marco Zompit et al., 2022). Collectively, these data demonstrate that the CIP2A-TOPBP1 complex responds robustly to diverse types of replication stress during mitosis, highlighting its broader role in the maintenance of genomic stability.

### CIP2A Mediates the Recruitment of MUS81, SLX4, and XPF, but Not RAD52, to Sites of Under-Replicated DNA in Mitosis

It was previously proposed that upon replication stress, TOPBP1 maintains genome integrity in mitosis by controlling chromatin recruitment of the structure-specific nuclease scaffold protein SLX4 (Pedersen et al., 2015). Since CIP2A acts as a regulator of TOPBP1 in mitosis, we considered the possibility that CIP2A may also be implicated in the recruitment of the SMX tri-nuclease complex.

We first examined FANCD2 recruitment, which classifies as an early upstream factor marking under-replicated DNA and facilitating the recruitment of key components that respond to replication stress (Chan et al., 2009). Notably, FANCD2 foci formation was unaffected by CIP2A depletion, indicating that CIP2A is not required for the initial recognition of under-replicated DNA (Fig. 2A). In addition, this suggests that replication stress at mitotic entry is comparable between control and CIP2A-depleted cells, ruling out the possibility that CIP2A deficiency reduces the DNA damage carried into mitosis.

**Figure 2:**
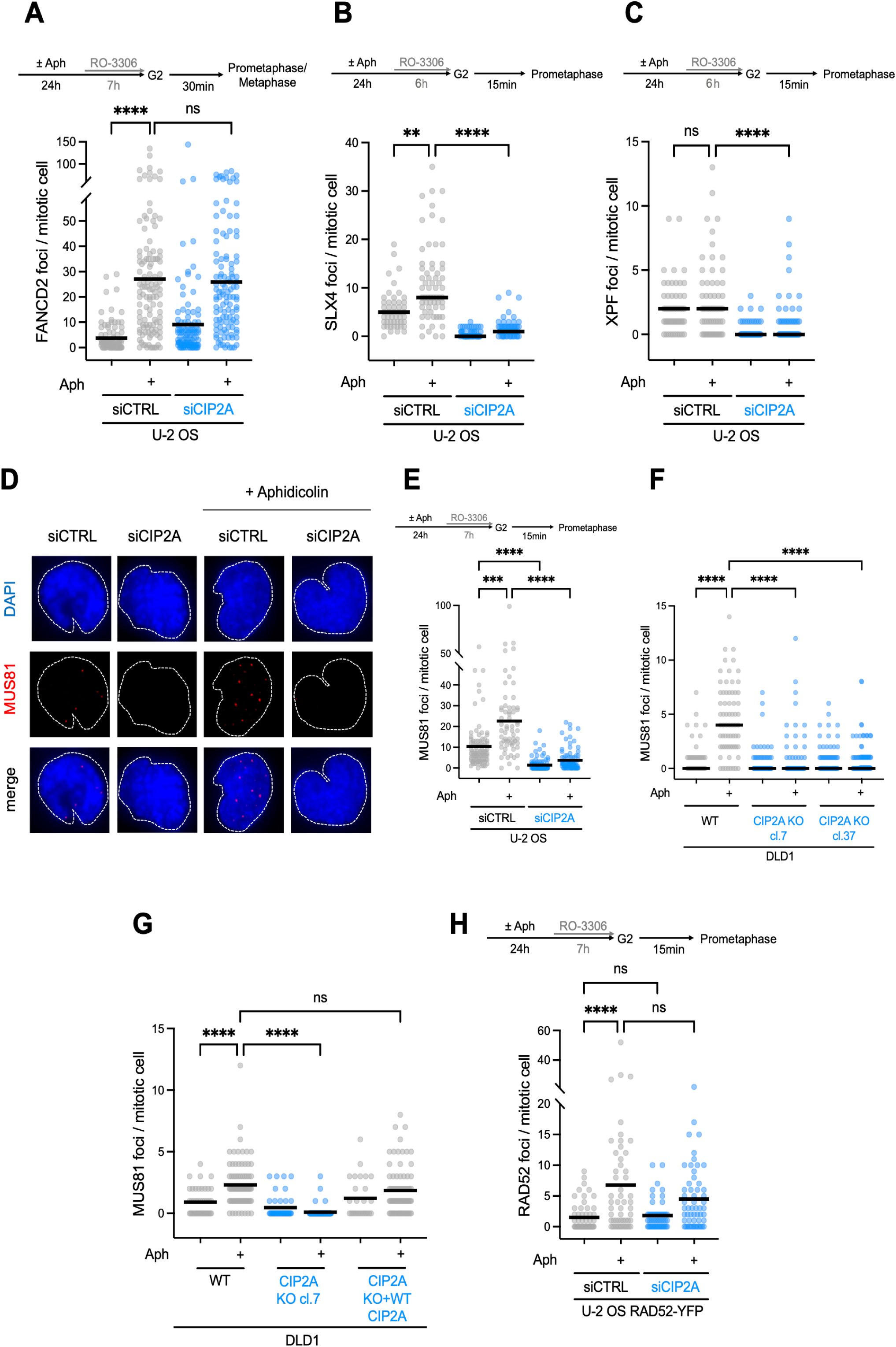
CIP2A mediates the recruitment of the SMX tri-nuclease complex, but not RAD52, to under-replicated DNA in mitosis. **A)** Dot plot showing FANCD2 foci per mitotic cell in U2OS cells treated as illustrated in the above panel, with median indicated. n = 3 independent replicates. **B)** Dot plot showing SLX4 foci per mitotic cell in U2OS cells treated as illustrated in the above panel. Median indicated. n = 2 independent replicates. **C)** Dot plot showing XPF foci per mitotic cell in U2OS cells treated as illustrated in the above panel. Median indicated. n = 2 independent replicates. **D)** Confocal microscopy images showing MUS81 foci (red) in mitotic U2OS cells, with nuclei stained with DAPI (blue). Images are maximum projections of Z-stacks. **E)** Dot plot showing MUS81 foci per mitotic cell in U2OS cells treated as illustrated in the above panel, with median indicated. n = 2 independent replicates. **F)** Dot plot showing MUS81 foci per mitotic cell in DLD1 cells treated as illustrated in panel depicted in **E)**, with median indicated. n = 2 independent replicates. **G)** Dot plot showing MUS81 foci per mitotic cell in DLD1 cells treated as in panel depicted in **E)**. Mean indicated. n = 4 independent replicates. **H)** Dot plot showing RAD52 foci per mitotic cell in U2OS cells treated as illustrated in the above panel, with median indicated. n = 3 independent replicates. Statistical analysis in all dot plot panels: Kruskal-Wallis test, followed by uncorrected Dunn’s multiple comparisons test.

Next, we analyzed the recruitment of SLX4, a scaffold protein that coordinates the activity and the recruitment of multiple structure-specific nucleases, among which MUS81-EME1 and XPF-ERCC1, thereby facilitating their accumulation at sites of DNA damage and under-replicated regions (Castor et al., 2013; Garner et al., 2013; Falquet et al., 2019). Replication stress induction strongly increased SLX4 foci formation in mitotic control cells, consistent with its involvement in MiDAS; however, depletion of CIP2A significantly decreased SLX4 foci number (Fig. 2B). In line with SLX4 function in recruiting XPF-ERCC1, we found that XPF foci were reduced in U2OS cells depleted of CIP2A (Fig. 2C). Similarly, we observed a significant decrease in MUS81 foci formation upon depletion of CIP2A in U2OS cells and in DLD1 CIP2A KO cells, respectively (Fig. 2D, E, F). Significantly, we observed a full rescue of MUS81 foci formation in DLD1 CIP2A KO cells expressing wild-type CIP2A (Fig. 2G), thus indicating that the observed MUS81 recruitment defect is not a clonal effect or caused by an off-target effect of the CIP2A siRNA.

Finally, we examined the recruitment of RAD52, which has also been implicated in the mitotic response to replication stress as a strand-annealing factor necessary to achieve BIR-like replication (Bhowmick et al., 2016). Unlike MUS81, XPF, and SLX4, RAD52 foci formation was unaffected by CIP2A depletion, suggesting that CIP2A selectively regulates the SLX4-MUS81-XPF recruitment without impacting RAD52 binding (Fig. 2H).

Collectively, these findings demonstrate that CIP2A is required for the recruitment of SLX4, XPF, and MUS81 to sites of under-replicated DNA during mitosis, most likely ensuring proper replication intermediate resolution, while having no impact on RAD52 recruitment.

### CIP2A Facilitates MiDAS in Some Cell Lines, but Only Partially in Others

Mitotic DNA synthesis (MiDAS) is a well-established mechanism that allows cells to cope with replication stress in mitosis, ensuring proper chromosome segregation (Bhowmick et al., 2023). It plays a crucial role in completing under-replicated DNA, thereby preventing chromosome mis-segregation and genomic instability. It was also proposed that MiDAS depends on the structure-specific nuclease MUS81 and its scaffold protein SLX4 (Minocherhomji et al., 2015). To determine whether CIP2A is involved in the MiDAS process, we monitored MiDAS activity in BRCA2-deficient cells, which exhibit spontaneous low levels of MiDAS activity (Feng and Jasin, 2017), and after treatment with replication stress-inducing agents, specifically ATR inhibition and Aph treatment. We first examined MiDAS in BRCA2 KO DLD1 cells by analyzing EdU incorporation on chromosome spreads from metaphase cells, which would provide high sensitivity to detect even low levels of MiDAS. Cells were synchronized at the G2/M boundary using the CDK1 inhibitor RO-3306, released into mitosis in the presence of EdU to label any DNA synthesis, and subsequently arrested in metaphase with Colcemid for chromosome spreading. Metaphase spreads were then analyzed for the number of EdU foci per chromosome. Consistent with a recent pre-print, we observed that BRCA2 KO cells exhibited increased EdU incorporation compared to WT cells, and that siRNA-mediated depletion of CIP2A reduced the number of EdU foci per chromosome (Fig. 3A) (Martin et al., 2024).

**Figure 3:**
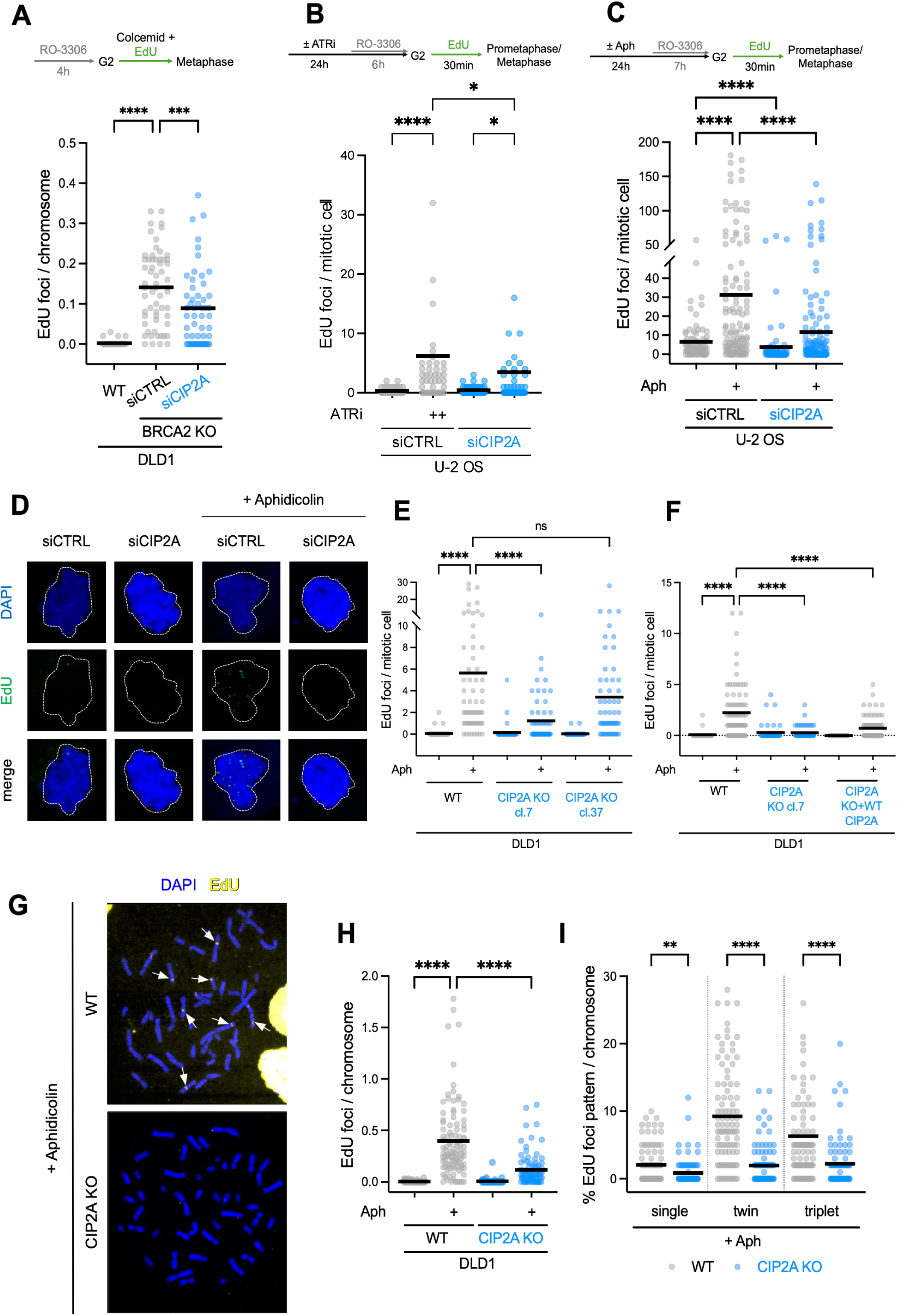
CIP2A involvement in MiDAS is cell line dependent. **A)** Dot plot showing EdU foci per chromosome in DLD1 cells, with each dot representing a single metaphase spread. Cells were synchronized with RO-3306 for 4 hours, then released in presence of Colcemid and EdU. Median indicated. n = 2 independent replicates. **B)** Dot plot showing EdU foci per mitotic cell in DLD1 cells treated as illustrated in the above depicted panel. Mean indicated. n = 2 independent replicates. **C)** Dot plot showing EdU foci per mitotic cell in U2OS cells treated as illustrated in the above depicted panel. Median indicated. n = 3 independent replicates. **D)** Confocal microscopy images showing EdU foci (green) in mitotic U2OS cells, with nuclei stained with DAPI (blue). Images are maximum projections of Z-stacks. **E)** Dot plot showing EdU foci per mitotic cell in DLD1 cells treated as in panel depicted in **C)**, with median indicated. n = 2 independent replicates. **F)** Dot plot showing EdU foci per mitotic cell in DLD1 cells including DLD1 CIP2A KO complemented with CIP2A WT, treated as in panel depicted in **C)**. Mean indicated. n = 2 independent replicates. **G)** Widefield microscopy images showing EdU foci (yellow) on DLD1 WT and CIP2A knockout metaphase spreads stained with DAPI (blue). White arrows indicate some representative EdU foci. **H)** Dot plot showing EdU foci per chromosome in DLD1 cells, with each dot representing a single metaphase spread. Cells were treated ± Aph for 24 hours and synchronized with RO-3306 during the last 4 hours, then released in presence of Colcemid and EdU for 40 min. Median indicated. n = 3 independent replicates. **I)** Quantitative analysis of EdU foci pattern in metaphase spreads shown in **G)**. EdU foci were classified as single, twin, or triplet. n = 3 indpendent replicates. Statistical analysis in all dot plot panels: Kruskal-Wallis test, followed by uncorrected Dunn’s multiple comparisons test.

Next, we investigated whether CIP2A contributes to MiDAS following ATR inhibition. U2OS cells were treated with ATRi, synchronized in G2/M, and released into mitosis in the presence of EdU, followed by immunofluorescence and quantification of mitotic EdU foci. Notably, cells exhibited a significant increase in mitotic EdU incorporation upon ATR inhibition, indicating that in unperturbed conditions, ATR suppresses MiDAS. We observed that CIP2A depletion is associated with a mild reduction of ATRi-induced MiDAS, further suggesting its involvement in the mitotic replication stress response (Fig. 3B).

MiDAS has been well characterized as a mechanism to complete replication of under-replicated DNA induced by Aph (Minocherhomji et al., 2015). To assess whether CIP2A is also required for Aph-induced MiDAS, we repeated these assays in U2OS and DLD1 cells, in which CIP2A was either depleted or knocked out, respectively. According to previous findings, U2OS cells exhibited high levels of MiDAS. Aph-induced MiDAS was strongly reduced upon CIP2A depletion in U2OS cells, as measured by immunofluorescence quantification of mitotic EdU foci (Fig. 3C, D). Surprisingly though, we observed generally low MiDAS activity in DLD1 wild type cells and a significant reduction of EdU foci only in one of our two previously published DLD1 CIP2A KO clones (Adam et al., 2021) (Fig. 3E), thus suggesting that CIP2A may not retain a crucial role in MiDAS regulation in DLD1 cells and that the MiDAS defect observed in one of the DLD1 CIP2A KO clones is a clonal effect, which is only partially caused by CIP2A loss. Supporting this notion, the re-expression of wild type CIP2A in this DLD1 CIP2A KO clone only minimally rescues the MiDAS defect (Fig. 3F).

Although aware of the discrepancy observed between the two DLD1 CIP2A KO clones regarding the role of CIP2A in mediating MiDAS, we sought to investigate whether CIP2A loss correlates with decreased EdU incorporation in metaphase spreads. To this end, we considered the DLD1 CIP2A KO clone 7, which showed clear diminished MiDAS activity upon CIP2A loss. We found a significant reduction of EdU incorporation per chromosome (Fig. 3G, H). Analysis of MiDAS foci patterns further revealed that CIP2A depletion preferentially induces twin and triplet EdU foci (Fig. 3I).

Altogether, these findings demonstrate that the impact of CIP2A loss on MiDAS is at least in part context dependent, which is consistent with findings reported in a recent pre-print (De Haan et al., 2025). Indeed, while CIP2A emerges as a key mediator of MiDAS across different replication stress conditions in U2OS cells, its involvement in promoting MiDAS in DLD1 cells appears more ambiguous. A possible rationale for this cell line-based discrepancy might be that U2OS cells rely on the alternative lengthening of telomeres (ALT) mechanism for the elongation of telomeres, in contrast to DLD1 cells, which express the telomerase enzyme. Specifically, ALT cells exhibit enhanced levels of telomere replication stress due to atypical telomeric sequences, dysregulation of telomeric chromatin, and altered nucleoprotein structure (Lu & Pickett., 2022). As a direct consequence of the heightened replication stress, it is tempting to speculate that ALT cells might be particularly dependent on MiDAS to complete the replication, and because of this, the role of CIP2A in promoting MiDAS might become more critical in these cells.

### Disruption of the CIP2A-TOPBP1 Interaction Abrogates MUS81 Recruitment But does Not Affect MiDAS

Our data suggest that MiDAS and SMX recruitment are not functionally linked, as CIP2A complementation in CIP2A KO cells does not correlate with a complete restoration of the MiDAS activity while rescuing MUS81 foci formation upon replication stress (Fig. 2G, Fig. 3F). To further explore the role of CIP2A in response replication stress, we sought to determine whether the disruption of the CIP2A-TOPBP1 interaction affects MiDAS activity and MUS81 foci formation. To this end, we used a previously described approach consisting of a DLD1 WT cell line expressing a fragment of TOPBP1, referred to as BRCT6-long (B6L, residues 756-1000) (Adam et al., 2021). This fragment is under the control of an FKBP12-derived destabilization domain and its expression, which is induced upon the treatment with Shield-1 (S1) ligand, associates with the disruption of the TOPBP1-CIP2A interaction. To confirm the abrogation of the interaction between CIP2A and TOPBP1, we performed a proximity ligation assay (PLA) in cells treated with Aph and/or Shield1 and observed a decrease in the mean intensity of the PLA signal for CIP2A and TOPBP1 upon the disruption of the CIP2A-TOPBP1 interaction (Fig. 4A, B). Supportive to this finding, DLD1 WT B6L expressing cells exhibit a reduction in both CIP2A and TOPBP1 foci following the treatment with Shield-1, independently of the Aph treatment (Fig. 4C, D). To determine the impact of the abrogation of the CIP2A-TOPBP1 interaction on MiDAS, DLD1 WT B6L expressing cells were synchronized in G2/M and released in the presence of EdU. In line with our previous conclusion that CIP2A is at least partially dispensable for MiDAS in DLD1 cells (Fig. 3F), we did not observe a significant reduction in EdU foci in Aph-stressed cells in which the expression of the B6L fragment was induced (Fig. 4E, F). However, MUS81 foci formation was significantly reduced in response to the disruption of the CIP2A-TOPBP1 interaction (Fig. 4G, H), thus providing additional confirmation that the recruitment of the SMX tri-nuclease complex and MiDAS do not correlate. Nevertheless, we cannot exclude the possibility that MUS81 is involved in the process of MiDAS. Although MUS81 accumulation into foci does not seem to be required for MiDAS, its role in cleaving under-replicated DNA regions might still occur through the recruitment of very few MUS81 molecules that may escape microscopic detection.

**Figure 4:**
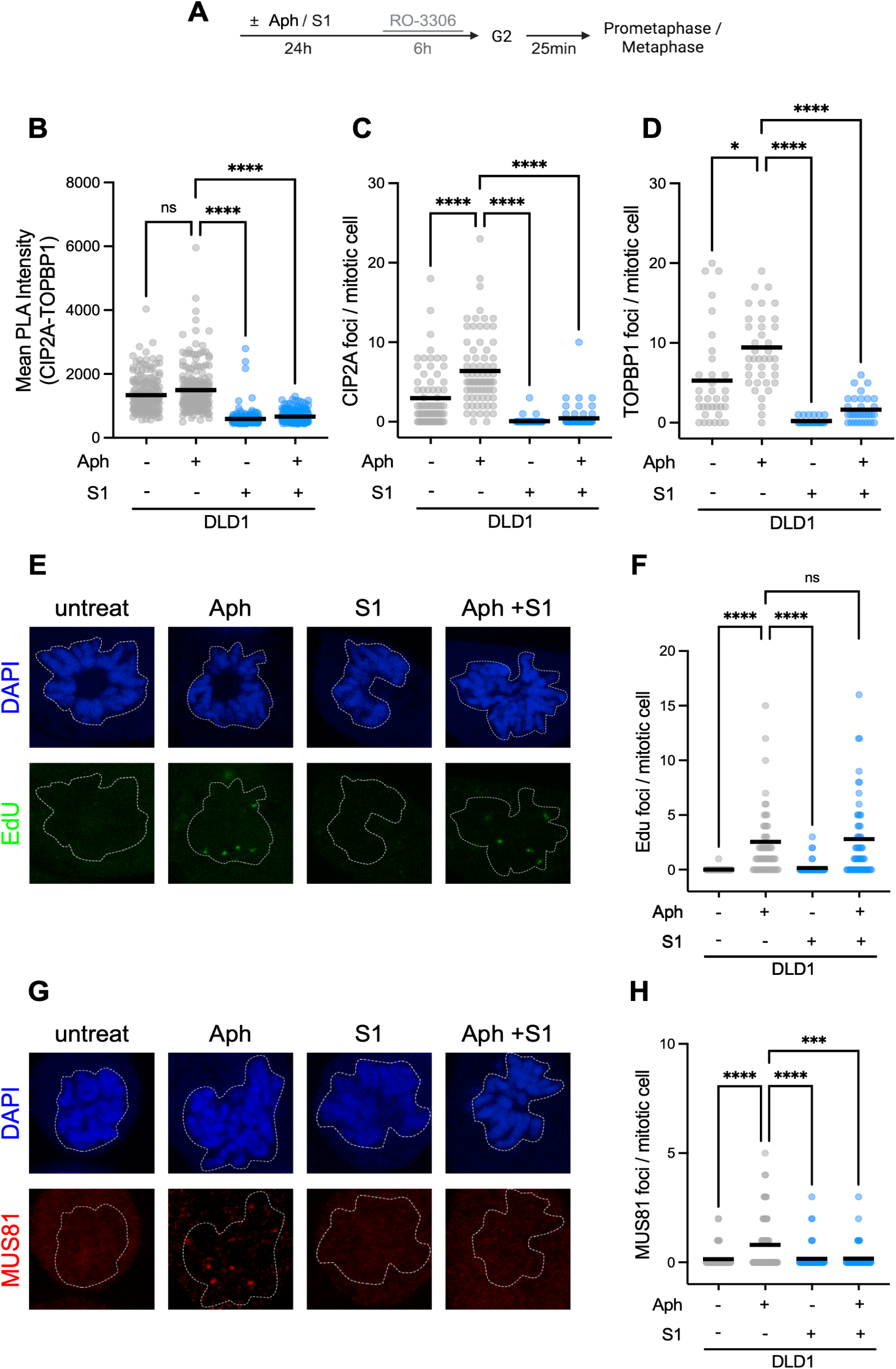
MUS81 recruitment, but not MiDAS, is affected by the disruption of CIP2A-TOPBP1 interaction. **A)** Schematic representation of the experimental workflow. **B)** Quantitative analysis of the mean proximity ligation assay (PLA) intensity in DLD1 WT B6L expressing cells treated as in **A)** and stained for CIP2A and TOPBP1. Statistical analysis: Brown-Forsythe and Welch ANOVA test. n = 2 independent replicates. **C)** Dot plot showing CIP2A foci per mitotic cell in DLD1 WT B6L expressing cells treated as in **A)**, with mean indicated. Cells were incubated with Shield-1 (S1) during release. Statistical analysis: Kruskal-Wallis test, followed by uncorrected Dunn’s multiple comparisons test. n = 2 independent replicates. **D)** Dot plot showing TOPBP1 foci per mitotic cell in DLD1 WT B6L expressing cells treated as in **A)**, with mean indicated. Cells were incubated with Shield-1 (S1) during release. Statistical analysis: Kruskal-Wallis test, followed by uncorrected Dunn’s multiple comparisons test. n = 2 independent replicates. **E)** Confocal microscopy images showing EdU foci (green) in mitotic DLD1 WT B6L expressing cells, with nuclei stained with DAPI (blue). Images are maximum projections of Z-stacks. **F)** Dot plot showing EdU foci per mitotic cell in DLD1 WT B6L expressing cells treated as in **A)**. Cells were incubated with EdU and Shield-1 (S1) during release. Mean indicated. Statistical analysis: Kruskal-Wallis test, followed by uncorrected Dunn’s multiple comparisons test. n = 2 independent replicates. **G)** Confocal microscopy images showing MUS81 foci (red) in mitotic DLD1 WT B6L expressing cells, with nuclei stained with DAPI (blue). Images are maximum projections of Z-stacks. **H)** Dot plot showing MUS81 foci per mitotic cell in DLD1 WT B6L expressing cells treated as in **A)**. Cells were incubated with Shield-1 (S1) during release. Mean indicated. Statistical analysis: Kruskal-Wallis test, followed by uncorrected Dunn’s multiple comparisons test. n = 2 independent replicates.

### CIP2A Loss Leads to Chromosomal Instability Upon Replication Stress

To investigate the consequences of CIP2A loss under replication stress, we analyzed genome instability markers following Aph treatment. Specifically, we examined anaphase defects and micronuclei formation, two hallmarks of chromosomal mis-segregation and incomplete DNA replication.

We first assessed lagging chromosome fragments in anaphase cells, which were collected 1 hour after release from G2/M synchronization. Control cells exhibited an increased frequency of lagging fragments upon treatment with a low dose of Aph. However, in CIP2A-depleted cells, Aph treatment led to a further significant increase in anaphase cells displaying lagging DNA fragments, indicating a failure to properly resolve DNA lesions caused by replication stress before chromosome segregation (Fig. 5A, B).

**Figure 5:**
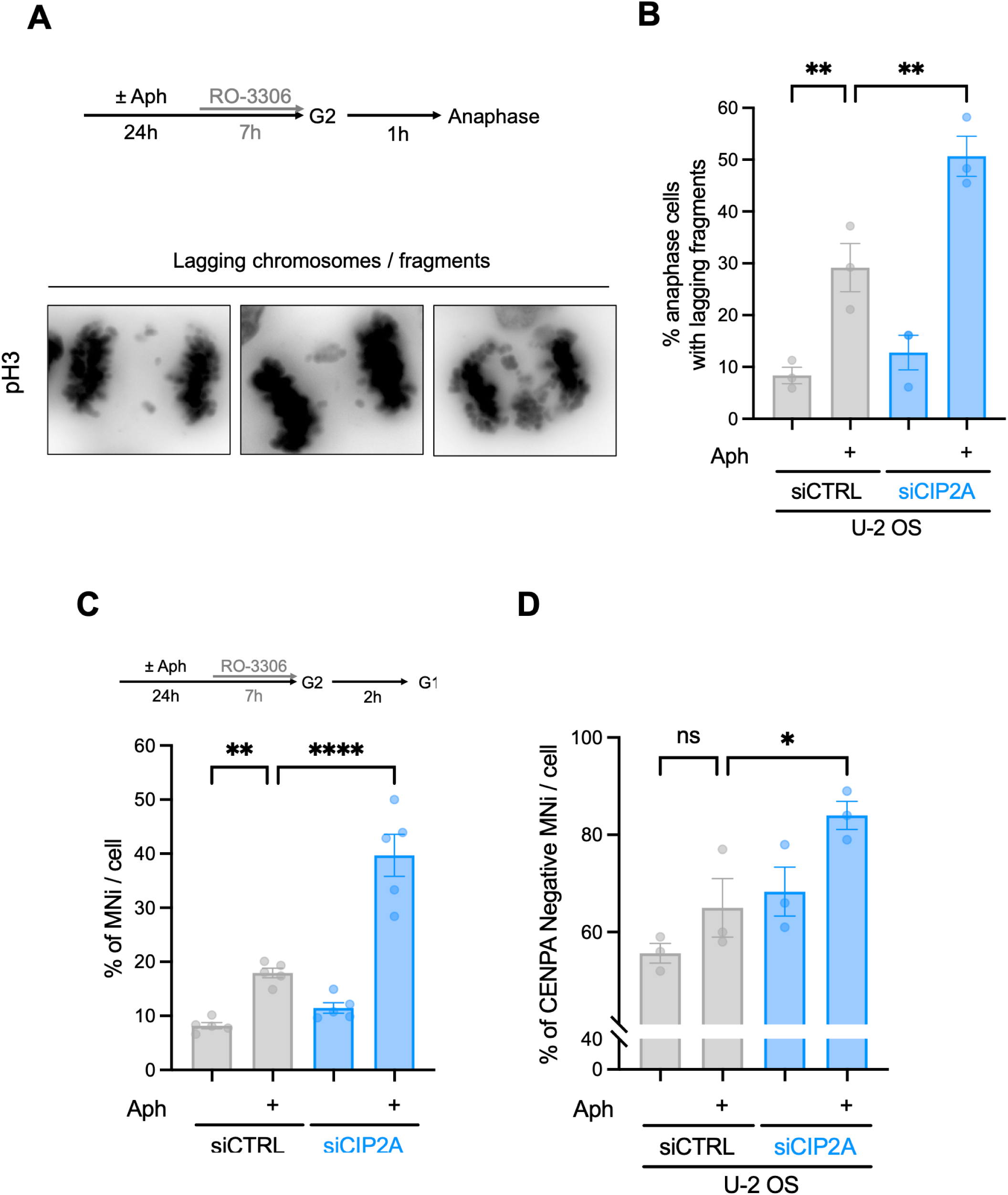
CIP2A loss is associated with chromo-somal instability in respo-nse to replication stress. **A)** Representative examples of anaphase cells with lagging chro-mosomes or chromosome fragments, stained with pH3. The experimental scheme used to collect anaphase cells is depicted. **B)** Mean ± SEM of anaphase U2OS cells displaying lagging chromosomes or chromo-some fragments. Statistical analysis: one-way ANOVA. n = 3 indpendent replicates. **C)** Mean ± SEM of micronuclei (MNi) per G1 cell. Statistical analysis: one-way ANOVA. n = 5 independent replicates. **D)** Mean ± SEM of CENP-A Negative micronuclei (MNi) per G1 cell. Statistical analysis: one-way ANOVA. n = 3 independent replicates.

Next, we examined micronuclei formation in G1 cells, which were collected 2 hours post-release from RO-3306 synchronization. CIP2A-depleted cells displayed a higher frequency of micronuclei compared to controls, suggesting that unresolved replication intermediates persist into the next cell cycle, leading to micronuclei containing the missegregated DNA fragments observed in anaphase cells (Fig. 5C). Finally, to determine whether these micronuclei resulted from broken chromosome fragments rather than entire mis-segregated chromosomes, we performed CENP-A staining. CENP-A-negative micronuclei were significantly enriched in CIP2A-depleted cells, confirming that these structures originated from acentric chromosome fragments (Fig. 5D).

Altogether, these results demonstrate that CIP2A is essential for ensuring the proper resolution of replication stress before mitotic exit, as its depletion leads to increased chromosomal mis-segregation and micronuclei formation following Aph treatment, ultimately compromising genome stability. In light of this, targeting CIP2A might prove a relevant therapeutic strategy for cancer characterized by significant levels of replication stress, in particular BRCA-deficient tumors. Indeed, the increased sensitivity of cells with enhanced levels of replication stress to CIP2A loss may make CIP2A a good drug target in cancers characterized by high levels of replication stress.

## Materials and methods

### Cell lines

Unless otherwise stated, all cells were cultured at 37 °C in a humidified incubator with 5% CO₂. DLD1 cells were maintained in RPMI-1640 medium supplemented with 10% fetal bovine serum (FBS) and 1% penicillin-streptomycin. DLD1 BRCA2 knockout (KO) cells were obtained from ATCC (HTB-96). DLD1 CIP2A KO clones 7 and 37 and DLD1 B6L expressing cells were a generous gift from Dan Durocher (Adam et al., 2021). U2OS were cultured in DMEM supplemented with 10% FBS and 1% penicillin-streptomycin. The U2OS YFP-RAD52 expressing cell line was a gift from Jiri Lukas (Bekker-Jensen et al., 2006). All cell lines were regularly tested for mycoplasma contamination.

### Gene silencing with RNA interference

Transient gene depletion was performed using Lipofectamine™ RNAiMAX Transfection Reagent (Invitrogen, 13778150), according to the manufacturer’s instructions. The following siRNAs are used at a final concentration of 20 nM: ON-TARGETplus Human KIAA1524 (57650) siRNA SMARTpool L-014135-01-0005 for CIP2A. The control siRNA (siCTRL, UGGUUUACAUGUCGACUAA-dTdT) is from obtained from Microsynth AG. Experiments were performed at 48-72 hours post-transfection.

### Drugs

The following compounds were utilized in this study at the following concentration, unless otherwise indicated in the corresponding figure legends: 0.4 µM Aph (MedChem Express, HY-N6733), 7 µM RO-3306 (SelleckChem, S7747), 0.1 µg/mL Karyomax Colcemid (Gibco, 1521201), 10 µM VE-821 ATR inhibitor (SelleckChem, S8007), 1 µM Shield-1 (Takara Bio, 632189).

### Immunofluorescence

Cells were grown on glass coverslips and fixed with 10% formalin (corresponding to 4% paraformaldehyde) for 15 min at room temperature (RT) or with ice-cold methanol for 15 min on ice. For SLX4 staining, pre-extraction with cold CSK buffer (100 mM PIPES pH 7, 100 mM NaCl, 300 mM sucrose, 3 mM MgCl_2_) supplemented with 0.5% Triton X-100 was performed for 8 min prior to fixation. For standard immunofluorescence staining, cells were permeabilized with 0.5% Triton X-100 in PBS for 15 min, then blocked with 10% FBS for 1 hour, followed by overnight incubation with primary antibodies diluted in 5% FBS. After three PBS washes, cells were incubated with Alexa Fluor-conjugated secondary antibodies (Thermo Fisher Scientific) at a 1:1000 dilution in 5% FBS for 1 hour. Following three additional PBS washes, coverslips were mounted on glass microscopy slides using VECTASHIELD® PLUS Antifade Mounting Medium containing 0.5 μg/mL DAPI (Vector Laboratories).

For MiDAS visualization, cells were permeabilized and blocked in 0.5% Triton X-100 with 3% BSA for 30 min. The Click-iT™ reaction (Thermo Fisher, Cat. No. C10337) was performed according to the manufacturer’s instructions, but using undiluted 10X additive as described (Garribba et al., 2018), for a total of 45 min. After a single wash with 3% BSA, primary antibodies were incubated overnight in 1% BSA, followed by the standard immunofluorescence protocol.

Images were acquired using a Zeiss LSM 900 confocal microscope with a 63x oil immersion objective. Mitotic cells were manually selected based on DAPI intensity and morphology. On average, 30–50 mitotic cells were imaged per condition using sequential scanning mode across a 6–8 μm z-stack with 8–12 optical sections.

### Image quantification

Total signal intensity and the number of foci per cell were quantified using customized pipelines developed in CellProfiler. Nuclei were segmented using the intensity-based Primary Object Detection module on the DAPI channel. Foci were identified either using the Primary Object Detection module or via the Find Maxima module, depending on the channel. The CellProfiler pipelines used in this study are available upon request. Micronuclei and anaphase cells with lagging chromosomes were quantified manually.

### Antibodies

The following primary antibodies were utilized in this study: anti-CIP2A (Mouse, Santa Cruz, sc-80659, 1/800), anti-TOPBP1 (Rabbit, Millipore, ABE1463, 1/250), anti-CENPA (Mouse, Abcam, ab13939, 1/500), anti-pSer10 Histone H3 (Rabbit, Millipore, 06-570, 1/1000), anti-MUS81 (Mouse, Santa Cruz, sc-53382, 1/250), anti-FANCD2 (Rabbit, Novus Biologicals, NB100-182, 1/200), anti-SLX4 (Sheep, generous gift from John Rouse (Wilson et al., 2013) 1/100), anti-XPF (Rabbit, Abcam, ab76948, 1/100). For GFP-tagged RAD52, coverslips were incubated with the ChromoTek GFP-Booster Alexa Fluor® 488 (Proteintech, gb2AF488-10, 1/500) for 1 h at RT.

### *In-situ* proximity ligation assay (PLA)

Cells were seeded on glass coverslips in 24-well plate at a density of 8 x 10^4^ cells per well and treated as indicated in the corresponding figure legends. Cells were synchronized in G2/M with 7 μM RO-3306 (Merck, 217699) 6 h before the end of the treatment. Mitotic release was performed by washing cells 3X with prewarmed PBS and by incubating the cells for 25 min in fresh medium. Coverslips were washed with cold PBS, fixed with ice-cold methanol for 10 min on ice and washed 3X with PBS. *In-situ* proximity ligation assay (PLA) was performed using Duolink^®^ PLA Reagents (Sigma-Aldrich, DUO92001, DUO92005, DUO92008, DUO82049) following the manufacture’s protocol. Cells were blocked for 1 h at 37 °C, followed by staining with anti-CIP2A (Mouse, Santa Cruz, sc-80659, 1:800) and anti-TOPBP1 (Rabbit, Millipore, ABE1463, 1:300) antibodies overnight at 4°C. Cells were then washed 2X with Wash Buffer A for 5 min and incubated with PLUS and MINUS PLA probes for 1 h at 37 °C. Cells were washed 2X with Wash Buffer A for 5 min. Ligation and polymerization steps were performed following the manufacturer’s instructions. Cells were washed 2X 10 min with Wash Buffer B, followed by a final wash with 0.01X Wash Buffer B. Coverslips were mounted on glass microscopy slides with VECTASHIELD^®^ PLUS Antifade Mounting Medium with DAPI. Quantification was performed with ImageJ (ROI Manager Tool, parameters: minimal and maximal gray value), following manual selection of mitotic cells.

### Metaphase Spreads

Cells were grown to 90% confluence on a 6 cm plate and were either treated or untreated with Aph (0.4 μM) for 24 hours and synchronized with RO-3306 (7 μM) for 4 hours. Cells were washed 3 x with pre-warmed PBS and released into EdU (40 μM) and KaryoMax Colcemid (0.1 μg/mL) for the last 40 min. Cells were trypsinized and transferred to a 15 ml Falcon™ tube, centrifuged at 176 g for 5 min and carefully resuspended in 2.5 ml of pre-warmed hypotonic buffer (15% FBS, 75 mM KCl) with intermittent agitation and incubated for 30 min at 37 °C. Cells were again pelleted at 176 g for 5 min, the supernatant was discarded, and the cell pellet was resuspended in 100 μl of hypotonic buffer. Cells were fixed by adding drop-wise 2.5 ml of ice-cold MeOH:AcOH 3:1 while slowly vortexing followed by 40 min on ice. After centrifugation at 176 g for 5 min at 4 °C, supernatant was discarded, and cells were resuspended in the remaining 70 μl of the fixation buffer. A total of 20μl of the cell suspension was then dropped at a 45° angle onto a wet glass slide and air-dried overnight.

Metaphases were stained using the Click-iT EdU Cell Proliferation Kit for Imaging, Alexa Fluor 488 dye (ThermoFisher Scientific). Firstly, slides were fixed with 4% formalin for 12 min, washed 3 x 5 min with PBS and blocked with 3% BSA for 30 min at RT. After permeabilization in 0.5% Triton-X in PBS for 20 min, slides were washed with PBS and a Click-iT reaction was performed according to the protocol provided by the supplier. Finally, slides were washed 3 x 5 min with 3% BSA in 0.5% Triton-X/PBS and metaphases were stained using VECTASHIELD® PLUS antifade mounting medium with DAPI and covered with a glass coverslip. Spreads were imaged using a Zeiss LSM 900 widefield microscope with a 40x water immersion objective. On average, 30–50 metaphase spreads were imaged per condition. Number of chromosomes per spread and EdU Foci were counted manually using ImageJ.

### Statistical analysis

The number of biological replicates is indicated in each figure legend as n = x. No statistical methods were used to predetermine sample size, and no inclusion or exclusion criteria were applied. Statistical analyses were performed using GraphPad Prism (version 10.4.1). The specific statistical test applied is indicated in the corresponding figure legends. Data are presented with the following significance thresholds: ns (not significant), P < 0.05 (*), P < 0.01 (**), P < 0.001 (***), and P < 0.0001 (****).

## Acknowledgements

We thank Daniel Durocher, Jiri Lukas and John Rouse for providing valuable reagents. We are grateful to Marcel ATM van Vugt for sharing unpublished results. Imaging was performed with equipment maintained by the Center for Microscopy and Image Analysis, University of Zürich. The Stucki lab is supported by grants from the Swiss National Foundation (310030_219462), Swiss Cancer Research (KFS-6185-08-2024) and by the Kanton of Zürich.

## Author contributions

The project was conceived and supervised by A.M. and M.S. Cell biological experiments were carried out and data was analyzed by A.M., A.P., A.H., N.V. and P.B-L. I.W. supervised P.B-L. Reagents were provided by A.A.S. The paper was written by A.M., A.P. and M.S. with contributions from the other authors.

